# Genetic variants affecting plant size and chemical defenses jointly shape herbivory in *Arabidopsis*

**DOI:** 10.1101/156299

**Authors:** AD Gloss, B Brachi, MJ Feldmann, SC Groen, C Bartoli, J Gouzy, ER LaPlante, CG Meyer, HS Pyon, SC Rogan, F Roux, J Bergelson, NK Whiteman

## Abstract

Herbivorous insects exhibit strong feeding preferences when choosing among plant genotypes, yet experiments to map loci mediating plant susceptibility to herbivory rarely incorporate host choice. To address this gap, we applied genome-wide association (GWA) mapping to uncover genetic polymorphisms mediating damage from foraging insects (two populations of *Scaptomyza flava*) across a mixture of *Arabidopsis thaliana* genotypes in experimental enclosures. The effect of chemical defenses (glucosinolates) on herbivory depended on herbivore genotype. Unlike many studies that minimize the effects of host choice behavior, we also found a large effect of plant size on herbivory—likely through its effect on plant apparency—that was independent of herbivore genotype. These herbivory-associated loci are polymorphic at fine spatial scales, and thus have potential to shape variation in herbivory within natural populations. We also show that the polymorphism with the largest effect on herbivory underlies adaptive latitudinal variation in *Arabidopsis* plant size across Europe. Overall, our results provide genetic support for ecological observations that variation in both chemical defenses and non-canonical defense traits (e.g., plant size and phenology) jointly shapes plant-herbivore interactions.

---

Interactions between plants and herbivores drive fundamental ecological and evolutionary processes. Herbivores remove 5-20% of the leaf tissue produced annually by plants ^1,2^, which reduces plant fitness ^3,4^, selects against susceptible plant genotypes ^4-6^, and in turn shapes the composition of ecological communities through effects that cascade across trophic levels ^7^. Understanding the mechanistic bases of plant-herbivore interactions is therefore a major goal in biology and agriculture ^8-10^.

Within plant species, individuals incur different amounts of damage from herbivores, and genetic differences among individuals explain a substantial proportion of this variation (e.g., ^11^). Identifying and characterizing specific genetic polymorphisms that shape herbivory-related phenotypes within plant populations, which we refer to as a gene-focused approach, offers a number of advantages for understanding how this variation arises and why it persists ^12-15^. First, in-depth functional studies of genes harboring these variants can uncover specific biochemical and physiological processes that mediate interactions with herbivores ^4^. Second, phenotypic comparisons among natural plant accessions or genetically engineered genotypes that vary at genes of interest can reveal how susceptibility to herbivory is linked to or trades off with other traits, including those of interest to evolutionary biologists (e.g., plant reproductive success ^16^) or crop breeders (e.g., plant biomass or yield). Third, population genetic studies, in which patterns at genes of interest are compared with neutral polymorphisms across the genome, can reveal how environmental pressures shape adaptive genetic variation within and among populations at loci affecting herbivory ^17^. Integrating these gene-focused methods provides a powerful approach to uncover traits and mechanisms that determine plant susceptibility to herbivory and to understand how these traits evolve ^12-15^. When conducted in crop species and their wild relatives, this approach can also find beneficial genetic variants that can be used to breed or engineer pest-resistant crops ^18^.

To develop a complete and unbiased understanding of the genetic basis of plant-herbivore interactions, a gene-focused approach should have the potential to identify genes and polymorphisms underlying any of the various strategies plants have evolved to mitigate herbivory. These strategies include evading, tolerating and resisting herbivores ^19,20^. In theory, genomic mapping techniques, such as genome-wide association studies (GWAS) and quantitative trait locus (QTL) analyses, are unbiased in this regard because they interrogate the entire genome for genetic polymorphisms that affect susceptibility to herbivory ^21,22^. In practice, however, the design of assays used to measure herbivory phenotypes often introduces bias by excluding the effects of herbivore foraging behavior. Many experiments place herbivores directly onto different plant genotypes or simulate herbivory (e.g., ^23-27^, but see ^28,29^). This approach yields precise measurements of plant resistance and herbivore performance phenotypes ^10^ and has enabled the discovery of novel loci shaping plant-herbivore interactions using genetic mapping techniques (e.g., ^23-27)^, but does so at the cost of minimizing the importance of whether a plant genotype will be encountered and attacked by a foraging herbivore. Consequently, the known catalog of genetic polymorphisms mediating plant-herbivore interactions is likely biased toward polymorphisms involved in resisting herbivory, such as those affecting physical and chemical defensive traits, while polymorphisms that mediate herbivore foraging behavior are likely underrepresented.

The fact that herbivore choice is often restricted in genetic mapping studies in the lab, whereas herbivores can move more freely in trait-based studies conducted in the field, may contribute to current uncertainty ^8^ regarding which plant traits are the primary determinants of herbivory. Co-evolutionary theory posits that plant defensive chemicals serve as the primary mechanism by which plants escape or reduce herbivory ^30-32^, and this prediction has empirical support from both trait-based ecological studies and genetic studies ^8,32^. The dominant role chemical defenses has been called into question by recent meta-analyses of trait-based ecological studies, which suggest that plant life history and morphological traits best explain differences in herbivory among plant genotypes or species, while effects of chemical defenses are smaller or context-dependent ^9,33,34^. One of the largest discrepancies between genetic mapping and trait-based studies specifically concerns the importance of plant traits (such as plant size) affecting probability that a plant will be encountered by herbivores, termed plant apparency ^20^. Plant apparency is correlated with susceptibility to herbivory in ecological studies (e.g., ^35-37),^ but because herbivore movement among plants is typically restricted or excluded in genetic mapping studies, loci affecting plant apparency have been conspicuously absent from the known catalog of genetic polymorphisms mediating plant-herbivore interactions. Mapping studies using experimental designs that incorporate insect foraging behavior, as a complement to precise assays focused on resistance phenotypes, could help reconcile results between genetic mapping and trait-based approaches.

Here, we dissected the genetic basis of susceptibility to herbivory in a genetic model plant, *Arabidopsis thaliana* (hereafter *Arabidopsis*), using a genetic mapping approach that simultaneously searched for loci involved in both resistance (for example, through chemical defenses) and evasion (for example, through reduced plant size) of herbivory. Adults of the herbivore species used in these assays—*Scaptomyza flava* (Diptera: Drosophilidae), a fly that naturally feeds on *Arabidopsis* in Europe and North America ^38^—frequently move among plants while foraging in nature. Adult *S. flava* females use a serrated ovipositor to puncture the undersides of leaves of plants in the family Brassicaceae ^38^, and they feed on sap that seeps into these wounds. Glucosinolates (GSLs), the major defensive chemicals in *Arabidopsis* ^39^, confer resistance to *S.flava* by slowing the feeding and growth rate of *S.flava* larval stages ^40^. However, the importance of intra-specific variation in plant GSL profiles for resistance to *S.flava* and the identity of other traits mediating resistance or evasion of herbivory *S. flava* are largely unknown.

To enable discovery of loci in the *Arabidopsis* genome involved in both evading and resisting herbivory by *S. flava*, we employed an experimental setup that allowed individual insects to forage among a genetically diverse set of plant genotypes and choose which genotypes they will attack. Our experimental design considered herbivory imposed by each of two populations of *S.flava*, one collected from Arizona (AZ) and another collected from Massachusetts (MA). We had three major goals: first, to identify genetic variants mediating susceptibility to foraging insect herbivores; second, to understand the physiological and ecological mechanisms through which these loci mediate plant-herbivore interactions; and third, to understand the evolution of these genetic variants in natural plant populations.

## RESULTS

### Plant genotype underlies variation in susceptibility to herbivory by foraging *Scaptomyza* flies

In controlled experiments where herbivores are placed directly onto individual plants, genetic differences among plants typically explain a substantial proportion of variation in herbivory (e.g., ^23^). If these genotypic effects are masked by stochasticity intrinsic to herbivore foraging decisions, experiments in which herbivores are allowed to forage may lack power to identify loci underlying variation in herbivory unless they employ extremely large sample sizes. To explore this problem, we investigated the feasibility of an experimental design that incorporates foraging behavior by allowing herbivores to potentially choose among all tested genotypes simultaneously (Fig. 1A-B). In each replicate, 288 natural *Arabidopsis* genotypes (accessions) from Europe were randomly arrayed along a grid within large enclosures in the laboratory (Fig. 1B), into which we introduced adult flies from either of two populations of *Scaptomyza flava*. Feeding punctures (“herbivory” hereafter) were quantified after 24 hours.

**Figure 1.**
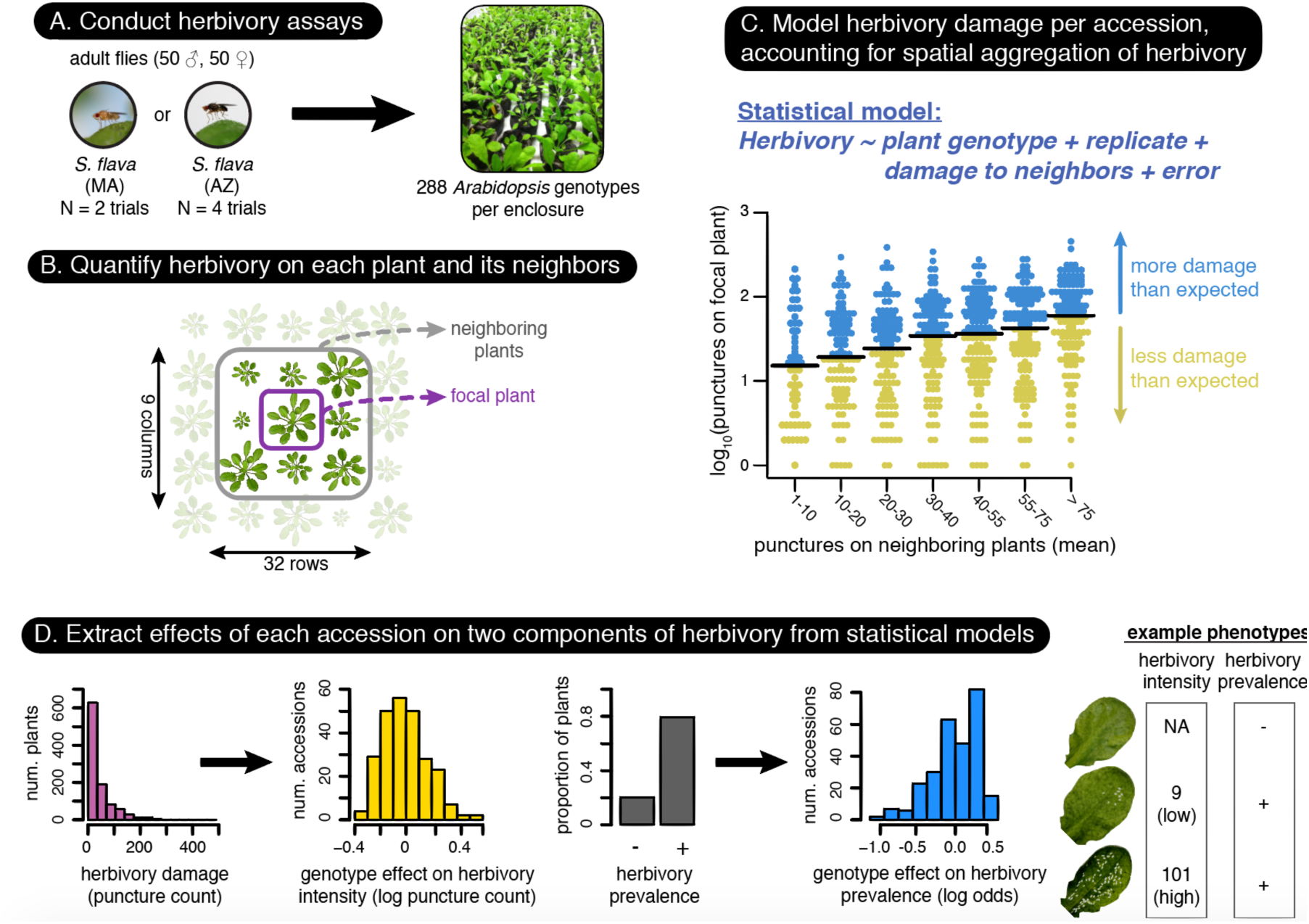
Experimental and statistical approach to quantify genetic variation in susceptibility to herbivory among *Arabidopsis* accessions. **(A)** We captured heritable variation in two metrics of herbivory damage—prevalence (presence/absence of herbivory per plant) and intensity (number of feeding punctures on damaged plants)—incurred when two populations of mustard-specialist *S. flava* flies foraged among a diverse mixture of plant genotypes. **(B)** Plants were arrayed in grids, with a single representative of each accession per experimental enclosure. **(C)** Generalized linear mixed models were used to estimate effect of accession genotype on herbivory damage phenotype, using the excess or deficit of herbivory relative to neighboring plants to account for stochastic spatial variation in herbivore foraging. **(D)** This approach yielded continuous distributions of heritable herbivory phenotypes across the *Arabidopsis* mapping population, shown here for herbivory by the *S. flava* AZ population.

We decomposed herbivory by *S. flava* into two components: (1) herbivory prevalence, defined as the presence or absence of herbivore damage per plant, which reflects host choice, and (2) herbivory intensity, defined as the amount of herbivore damage per damaged plant, which reflects feeding rate and/or duration. Herbivory prevalence and intensity were not correlated across *Arabidopsis* accessions (for assays with *S. flava* AZ: Spearman’s *ρ* = 0.02, *P* = 0.74; *S. flava* MA: *ρ* = -0.05, *P* = 0.55), and therefore reflect different phenotypes with different genetic bases.

Although plant accessions were randomly positioned in our experimental enclosures, herbivore damage was clustered spatially within each replicate (Fig. S1). Spatial clustering can arise stochastically when rates of host encounter and attack success are not completely deterministic ^41^. Failure to account for this aggregation increases uncertainty in measured phenotypes. Thus, we modeled herbivory phenotypes using an analytical framework that quantified the relative excess or deficit of herbivory damage per plant relative to neighboring plants (Fig. 1B-D). Compared to an approach that did not account for stochastic aggregation of herbivory, our approach improved the fit of the statistical model used to estimate herbivory phenotypes for each accession, increased the proportion of variation in herbivory explained by plant genotype (broad-sense heritability, *H^2^*), and increased power to identify putatively causal loci through GWA mapping (Table 1). Heritability estimates for herbivory phenotypes in this study were lower than typically observed for traits that do not involve biotic interactions ^42^, but fell within the range of estimates observed for herbivory phenotypes in other studies ^26^.

**Table 1.** Model fit (AIC), statistical significance of accession (i.e., genotype) effect, heritability, and statistical significance of the top trait-associated SNP from GWA mapping before and after accounting for spatial aggregation of herbivory within experimental assays.

| Trait | Without spatial analysis |  |  |  | With spatial analysis |  |  |  |  |
| --- | --- | --- | --- | --- | --- | --- | --- | --- | --- |
| | AIC | $P_{\text{Accession}}$ | $H^2, ^a$ | Best $P_{\text{SNP}}$ | AIC | $P_{\text{Accession}}$ | $H^2, ^a$ | $h^2, ^b$ | Best $P_{\text{SNP}}$ |
| Herbivory prevalence<br>(*S. flava* AZ) | 982.8 | $2.2 \times 10^{-2}$ | 0.10 | $1.3 \times 10^{-7}$ | 925.6 | $7.5 \times 10^{-3}$ | 0.14 | 0.13 | $1.9 \times 10^{-8}$ |
| Herbivory prevalence<br>(*S. flava* MA) | 561.4 | $1.2 \times 10^{-3}$ | 0.28 | $1.6 \times 10^{-6}$ | 521.1 | $1.7 \times 10^{-4}$ | 0.40 | 0.34 | $2.0 \times 10^{-7}$ |
| Herbivory intensity<br>(*S. flava* AZ) | 7836.4 | $3.6 \times 10^{-2}$ | N/A | $4.4 \times 10^{-6}$ | 7809.5 | $1.4 \times 10^{-3}$ | N/A | 0.57 | $1.9 \times 10^{-6}$ |
| Herbivory intensity<br>(*S. flava* MA) | 2744.9 | 0.10 | N/A | $8.7 \times 10^{-7}$ | 2736.1 | 0.10 | N/A | 0.51 | $4.1 \times 10^{-7}$ |

<sup>a</sup> Broad-sense heritability: the proportion of phenotypic variation explained by additive and non-additive genetic variation; estimated from variance components of GLMMs fitted to replicated phenotype measurements across accessions. Note: because herbivory intensity was modeled with a negative binomial error distribution, conventional methods to estimate heritability could not be applied.
<sup>b</sup> Narrow-sense heritability: the proportion of phenotypic variation explained by additive genetic variation; estimated as the proportion of phenotypic variation among accessions explained by pairwise relatedness coefficients (i.e., pseudo-heritability, as proposed in <sup>44</sup>). Note: the distribution of accession effects on herbivory phenotypes inferred using our non-spatial method was highly discontinuous, preventing estimation of $h^2$ .

### GWA mapping uncovers loci linking independent effects of plant size and chemical defenses to herbivory

To identify loci with large effects on herbivory, we conducted GWA mapping independently for each of four herbivory traits: prevalence and intensity of herbivory by *S. flava* AZ, and prevalence and intensity of herbivory by *S. flava* MA. Given the central importance of chemical defenses and plant size in ecological studies of plant herbivore interactions ^8^, ^9^, we also used GWA mapping to identify loci with effects on these traits (GSLs: aliphatic and indolic amino-acid derived GSLs; size: aboveground dry biomass, wet biomass, and rosette diameter) and treated these loci as *a priori* candidates affecting herbivory (Table S1). Our GWAS interrogated ~165,000 common SNPs distributed across the *Arabidopsis* genome and controlled for confounding due to population structure by including a matrix of relatedness among accessions as a random effect in a mixed model framework ^43,44^. With the exception of a previous dataset for aliphatic GSL profiles ^17^, all analyses relied on phenotypic measurements generated in the present study.

#### Effects of plant size and GSLs on herbivory

The strongest, statistically significant association with plant size phenotypes was restricted to a trio of SNPs in near-complete linkage disequilibrium in a narrow region of *Arabidopsis* chromosome 1 (Figs. 2A,S2 & Table 2). We refer to this genomic region, which had not previously been linked to variation in plant size, as the *Plant Biomass and Size Locus* (PBSL). We replicated the association between PBSL genotype and plant size through analysis of phenotypes from a previous GWAS for plant diameter (one-tailed *P* = 0.019; Fig. S3), which included some overlap with accessions used in our study but involved phenotypes generated independently in another growing location ^42^. Thus, the phenotypic effect of this locus is not unique to environmental conditions in our laboratory. Strikingly, the allele at PBSL associated with reduced plant size (i.e., narrower rosette diameter and smaller biomass) was also associated with reduced prevalence of herbivory by both herbivore populations (Fig. 2A & Table 2). This locus had a large effect, explaining over 10% of the variation in herbivory prevalence and plant size phenotypes in our mapping population (Table 2).

**Figure 2.**
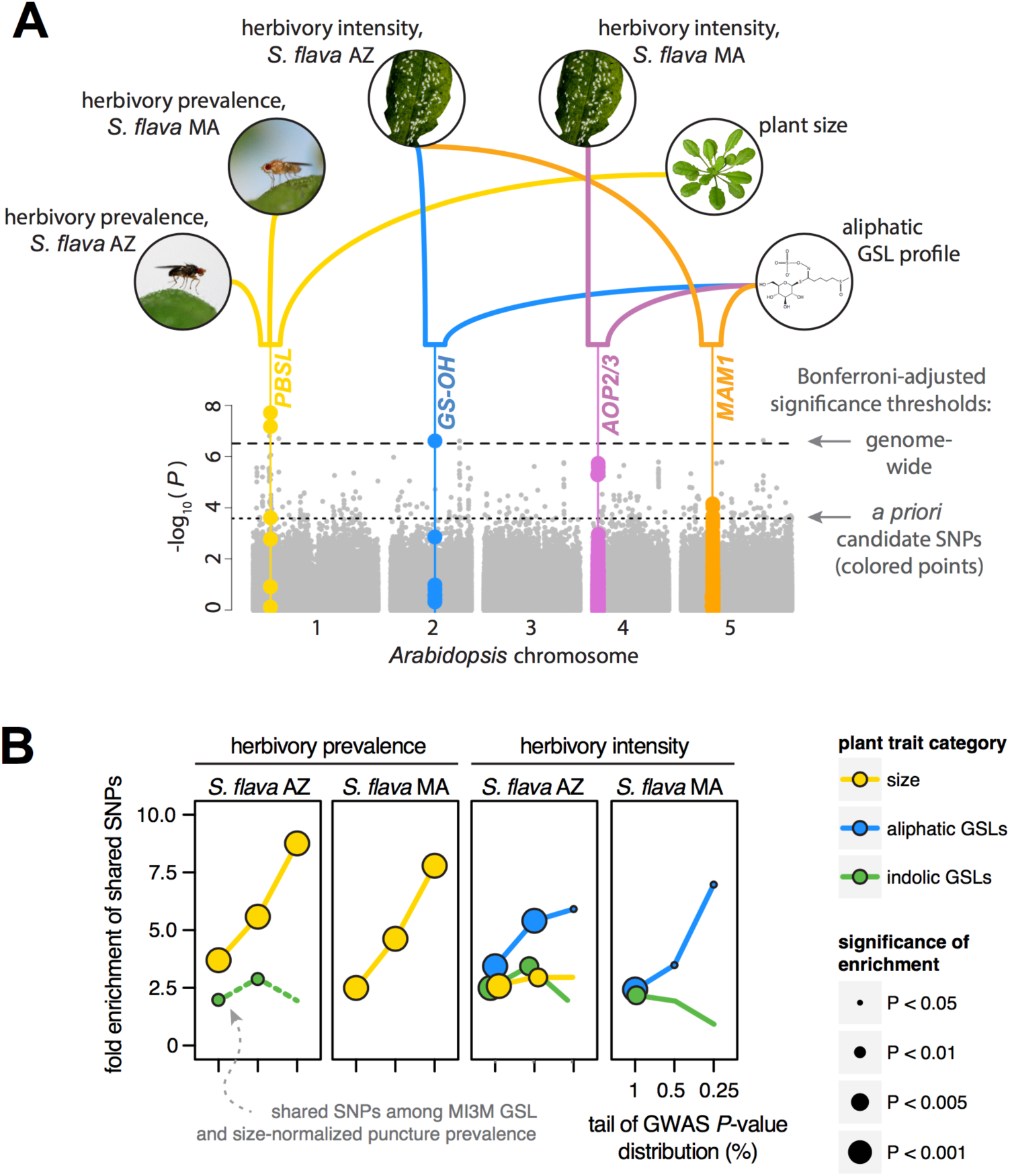
Genetic variation in plant size and GSL profile is associated with variation in herbivory by two herbivore populations. **(A)** Combined Manhattan plot for four herbivory phenotypes analyzed. SNPs associated with plant size (rosette mass or diameter) or GSL profile are colored and enlarged. Colored lines above the plot indicate associations with plant size, GSL, and herbivory traits. Significance was conservatively determined using a Bonferroni cutoff of *P* < 0.1, adjusting for the total number of SNPs analyzed (genomewide threshold) or the number of SNPs associated with plant size or GSLs (*a priori* candidate SNPs). **(B)** Fold enrichment of shared SNPs in the upper tail of GWAS *P-*value distributions for herbivory and either size or GSL phenotypes. Statistical significance was calculated using 1000 genomic permutations (see methods) and presented without correction for multiple tests. Only traits with a significant enrichment of shared SNPs in the 1% tail of the GWAS *P-*value distribution are shown (Bonferroni adjustment for 12 tests, *α* = 0.05), with the exception of a nominally significant (uncorrected *P* < 0.01) enrichment of indolic GSL-associated SNPs for herbivory prevalence by *S. flava* AZ after regressing out the effect of plant size prior to GWA mapping.

**Table 2.**
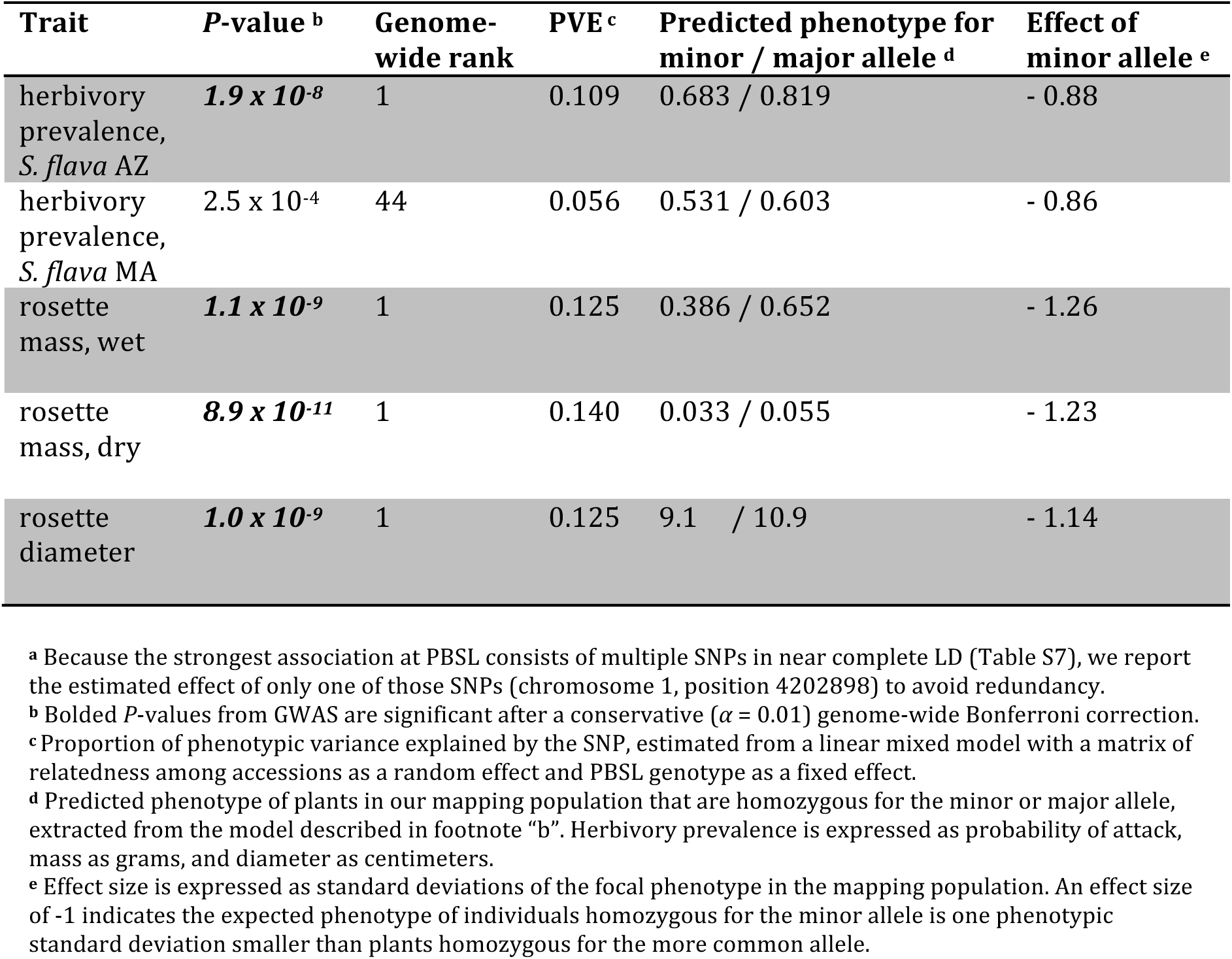
Effect of genetic variation at the *Plant Biomass and Size Locus* (PBSL)a on plant size and herbivory prevalence.

| Trait | *P*-value <sup>b</sup> | Genome-wide rank | PVE <sup>c</sup> | Predicted phenotype for minor / major allele <sup>d</sup> | Effect of minor allele <sup>e</sup> |
| --- | --- | --- | --- | --- | --- |
| herbivory prevalence, *S. flava* AZ | **$1.9 \times 10^{-8}$** | 1 | 0.109 | 0.683 / 0.819 | - 0.88 |
| herbivory prevalence, *S. flava* MA | $2.5 \times 10^{-4}$ | 44 | 0.056 | 0.531 / 0.603 | - 0.86 |
| rosette mass, wet | **$1.1 \times 10^{-9}$** | 1 | 0.125 | 0.386 / 0.652 | - 1.26 |
| rosette mass, dry | **$8.9 \times 10^{-11}$** | 1 | 0.140 | 0.033 / 0.055 | - 1.23 |
| rosette diameter | **$1.0 \times 10^{-9}$** | 1 | 0.125 | 9.1 / 10.9 | - 1.14 |

<sup>a</sup> Because the strongest association at PBSL consists of multiple SNPs in near complete LD (Table S7), we report the estimated effect of only one of those SNPs (chromosome 1, position 4202898) to avoid redundancy.
<sup>b</sup> Bolded *P*-values from GWAS are significant after a conservative ( $\alpha = 0.01$ ) genome-wide Bonferroni correction.
<sup>c</sup> Proportion of phenotypic variance explained by the SNP, estimated from a linear mixed model with a matrix of relatedness among accessions as a random effect and PBSL genotype as a fixed effect.
<sup>d</sup> Predicted phenotype of plants in our mapping population that are homozygous for the minor or major allele, extracted from the model described in footnote “b”. Herbivory prevalence is expressed as probability of attack, mass as grams, and diameter as centimeters.
<sup>e</sup> Effect size is expressed as standard deviations of the focal phenotype in the mapping population. An effect size of -1 indicates the expected phenotype of individuals homozygous for the minor allele is one phenotypic standard deviation smaller than plants homozygous for the more common allele.

Previous studies found that variation in the abundance of different aliphatic GSL compounds is primarily controlled by three loci, which are known to contain genes involved in GSL side chain modification ^17,39^ (Fig. S4). In our assays, GSL-associated SNPs at all three loci also significantly affected herbivory intensity (Fig. 2A), but the loci affecting herbivory by the *S. flava* MA population and the *S. flava* AZ population were distinct: the GS-OH and MAM loci affected herbivory by *S. flava* AZ, while the AOP locus affected herbivory by *S. flava* MA. Together, our GWA mapping results reveal a general effect (independent of herbivore genotype) of plant size and a context-dependent effect (dependent on herbivore genotype) of GSLs on herbivory.

Mapping studies in *Arabidopsis* have found that the tail of the GWAS *P*-value distribution is enriched for loci with true effects on plant phenotypes, even though many of these associations are not significant at a conservative genome-wide threshold ^42^. For each herbivory trait, we quantified fold enrichment of SNPs in the tail of the *P*-value distribution shared with plant size, aliphatic GSL profile, or indolic GSL profile. Results were consistent with our conservative single-locus approach (Fig. 2B). Top SNPs associated with herbivory prevalence by either herbivore population were strongly enriched for plant size-associated SNPs. In contrast, top SNPs associated with herbivory intensity by either herbivore population were strongly enriched for both aliphatic and indolic GSL-associated SNPs. We also detected weaker overlap between indolic GSLs and herbivory prevalence, and between plant size and herbivory intensity (despite no significant association at PBSL), but only for the *S. flava* AZ population.

#### Non-candidate traits affecting herbivory prevalence

To identify additional loci and traits that may shape patterns of herbivory in *Arabidopsis*, we focused on the four genomic regions that significantly affected herbivory prevalence but not plant size or GSL profile. These regions lacked obvious candidate genes, so we investigated whether any of these SNPs exhibited strong associations in GWAS of over 70 traits related to defense, development, and flowering time in *Arabidopsis* ^42^. Strikingly, the locus most strongly associated with herbivory prevalence (aside from PBSL) was consistently among the top genome-wide associations with plant age at the onset of flowering (Table S2). Because we excluded plants that had begun bolting from our analyses of herbivory, we hypothesize that variation at this locus may affect herbivory through physiological changes that precede the initiation of flowering. Although our subsequent analyses focus on the effects of plant size and chemical defenses, the putative effect of plant phenology on herbivory by *S. flava* warrants further study as another mechanism through which traits other than canonical defenses mediate susceptibility to herbivory ^9^ in *Arabidopsis.*

### Shared genetic architecture leads to correlations between herbivory and both plant size and chemical defense phenotypes

To complement inferences from GWAS, we used multiple regression to determine how heritable variation among accessions in plant size and GSL phenotypes relates to herbivory phenotypes. In addition to allowing the relative effects of each trait on herbivory to be estimated, a multiple regression approach might also identify relationships that were undetected through GWAS, given that GWAS has low power to map variants that are at low frequency or have small phenotypic effects. We focused on herbivory by the *S. flava* AZ population, where our herbivory assays had the most replication. For each herbivory trait, the regression model included the following phenotypes as putative predictors of herbivory: above-ground wet biomass (a component of plant size), the first two principal component (PC) axes that together describe ~80% of species-wide variation in the major aliphatic GSL compounds in *Arabidopsis* ^17^ (Fig. 3A), and the abundance of two indolic GSLs. Each model also included a kinship matrix to guard against spurious relationships that could arise if herbivory is affected by traits that were not included in our model and are geographically distributed similarly to GSLs or plant size phenotypes. Although plant size and some GSL traits were correlated, perhaps reflecting a tradeoff between growth and defense ^45^ or other sources of pleiotropy, these correlations are weak and should not prevent accurate estimation of the independent effects of GSLs and plant size on herbivory (Fig. S5).

**Figure 3.**
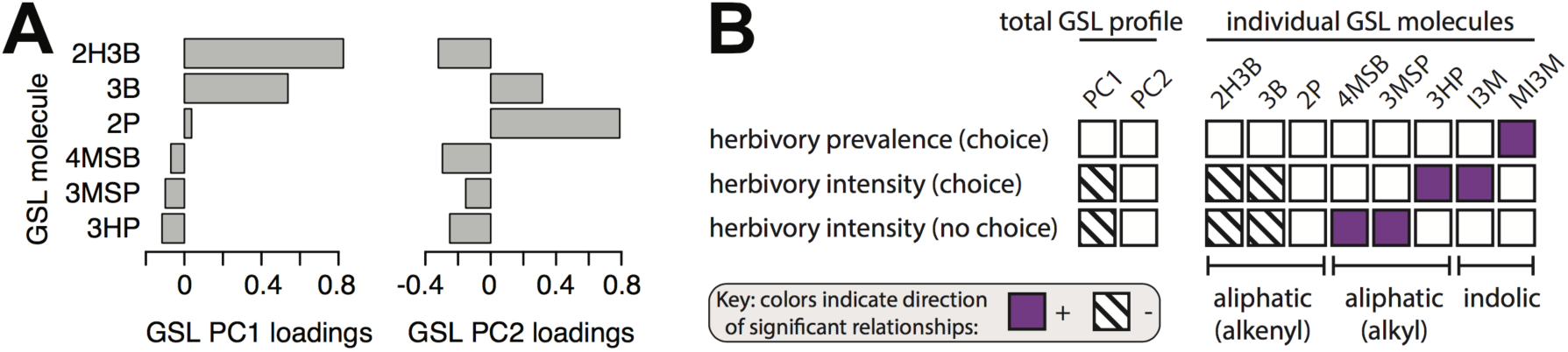
The effect of GSL profile varies among components of herbivory by *S. flava* AZ flies. **(A)** Contribution of six aliphatic GSLs to the first two principal components of GSL profile, which explain 80% of GSL variation among accessions (PC1: 50%, PC2: 30%). *N* = 585 accessions. **(B)** Effects of foliar abundances of different GSL molecules on variation in herbivory among accessions. Only effects that were significant at a 5% false discovery rate for 24 tests are shown. Because foliar abundances of some GSL molecules were strongly correlated, each model included only a single GSL molecule or PC as a predictor variable. To minimize the possibility of spuriously significant results caused by other geographically structured traits, each model also included plant size (wet mass) and a matrix of relatedness among accessions as predictor variables. *N* =182 accessions for herbivory prevalence, 164 accessions for herbivory intensity in choice assays, and 106 accessions for herbivory intensity in no choice assays. 2P: 2-propenyl; 2H3B: 2-hydroxy-3-butenyl; 3B: 3-butenyl; 3MSP: 3-methylsulfinylpropyl; 3HP: 3-hydroxypropyl; 4MSB: 4-methylsulfinylbutyl; I3M: indole-3-yl-methyl; MI3M: methoxy-indole-3-yl-methyl.

Our multiple regression approach revealed that plant size and GSL profiles together explained approximately 20% of the variation in susceptibility to herbivory among accessions (Table 3). However, their relative importance differed among herbivory traits. Plant size affected herbivory prevalence (*R^2^* = 14.5%) more than herbivory intensity (*R^2^* = 6.0%). In contrast, GSL phenotypes explained more variation in herbivory intensity (*R^2^* = 9.6%) than prevalence (*R^2^* = 3.2%). These results suggest that plant size has a greater effect than GSLs on host choice by *S. flava*, while plant size and GSLs are similarly strong determinants of feeding rate once a plant is attacked.

**Table 3.** Multiple regression models for the effects of plant size and GSLs on herbivory by *S. flava* AZ.

| Herbivory Phenotype | $\mu^a$ | N<sup>b</sup> | Effect of Plant Size: | | | Effect of GSLs: | | | | Effect of Plant Size and GSLs (full model<sup>e</sup>): | |
| --- | --- | --- | --- | --- | --- | --- | --- | --- | --- | --- | --- |
| | | | *P* | $\beta^c$ | *R*<sup>2</sup> | Phenotype<sup>d</sup> | *P* | $\beta^c$ | *R*<sup>2</sup> | *P* | *R*<sup>2</sup> |
| herbivory prevalence<br>(choice tests) | 1.37 | 182 | 1.1×10<sup>-8</sup> | 0.40 | 0.145 | PC1 | 0.67 |  |  | 9.7×10<sup>-9</sup> | 0.186 |
|  |  |  |  |  |  | PC2 | 0.98 |  |  |  |  |
|  |  |  |  |  |  | I3M | 0.38 |  |  |  |  |
|  |  |  |  |  |  | MI3M | 7.2×10<sup>-3</sup> | 0.19 | 0.032 |  |  |
| herbivory intensity<br>(choice tests) | 3.64 | 164 | 5.1×10<sup>-4</sup> | 0.13 | 0.060 | PC1 | 5.5×10<sup>-4</sup> | -0.15 | 0.061 | 4.5×10<sup>-8</sup> | 0.206 |
|  |  |  |  |  |  | PC2 | 0.23 |  |  |  |  |
|  |  |  |  |  |  | I3M | 4.6×10<sup>-4</sup> | 0.14 | 0.035 |  |  |
|  |  |  |  |  |  | MI3M | 0.70 |  |  |  |  |
| herbivory intensity<br>(no choice tests) | 40.4 | 106 | 4.6×10<sup>-4</sup> | 4.22 | 0.094 | PC1 | 0.02 | -2.58 | 0.040 | 5.5×10<sup>-5</sup> | 0.172 |
|  |  |  |  |  |  | PC2 | 0.65 |  |  |  |  |
|  |  |  |  |  |  | I3M | 0.41 |  |  |  |  |
|  |  |  |  |  |  | MI3M | 0.36 |  |  |  |  |

<sup>a</sup> Expected mean herbivory phenotype in the mapping population, estimated as the intercept from the mixed model used to quantify the effect of plant genotype (i.e., accession) on herbivory. Units are log odds for herbivory prevalence in choice tests, log puncture count for herbivory intensity in no choice tests, and puncture count for herbivory intensity in no choice tests.
<sup>b</sup> Number of accessions with available measurements for all predictor traits (plant size and GSL profile) and the response trait (a given herbivory phenotype) in each multiple regression model.
<sup>c</sup> Regression coefficients, scaled to indicate the expected difference in herbivory phenotype between a plant in the 10<sup>th</sup> percentile and 90<sup>th</sup> percentile of a given predictor trait in our mapping population.
<sup>d</sup> Four GSL phenotypes, including two principal components describing the abundance of 22 aliphatic GSLs (PC1, PC2) and abundances of two indolic GSLs (I3M, MI3M), were included in each model.
<sup>e</sup> Significance and *R*<sup>2</sup> were estimated for a “full model” that included only significant plant size and GSL phenotypes as predictive traits. Interaction terms among predictive traits were not significant (*P* > 0.05) and are not reported.

We also found that while GSL profile was associated with both herbivory prevalence and intensity, these two metrics of herbivory were affected by different GSL molecules. Herbivory prevalence was positively associated with the abundance of an indolic GSL (MI3M: methoxy-indole-3-yl-methyl). In contrast, herbivory intensity was negatively associated with the first PC axis describing GSLs (which primarily reflects an increased abundance of alkenyl GSLs with elongated side chain structures; Fig. 3A) and positively associated with the abundance of another indolic GSL (I3M: indole-3-yl-methyl). These results were robust to the different approaches to quantify GSL variation: models where each GSL compound was included separately, rather than integrated into PC axes, yielded consistent results (Fig. 3B). We also replicated the effect of alkenyl GSLs on herbivory intensity in additional experiments in which flies were not allowed to choose among accessions (“no choice” assays using a subset of the mapping population, Fig. 3B & Table 3), which further supports the conclusion that alkenyl GSLs impact *S. flava* feeding rate independently of host choice behavior.

### Assays with genetic mutant plants validate effects of plant size and chemical defenses on herbivory

#### Effect of GSLs

We validated the effects of GSL defenses on herbivory prevalence through additional assays using *Arabidopsis* mutants. In each assay, flies were allowed to forage among a mixture of two genotypes: plants with normal GSL profiles, or mutant plants ^46-48^ deficient in some or all GSL compounds. Consistent with our GWAS that explored natural genetic variation among accessions, these assays revealed that the effect of indolic GSLs on host preference is dependent on herbivore genotype: indolic GSLs were attractive to *S. flava* AZ flies, but had no effect on foraging by *S. flava* MA flies (Fig. 4A).

**Figure 4.**
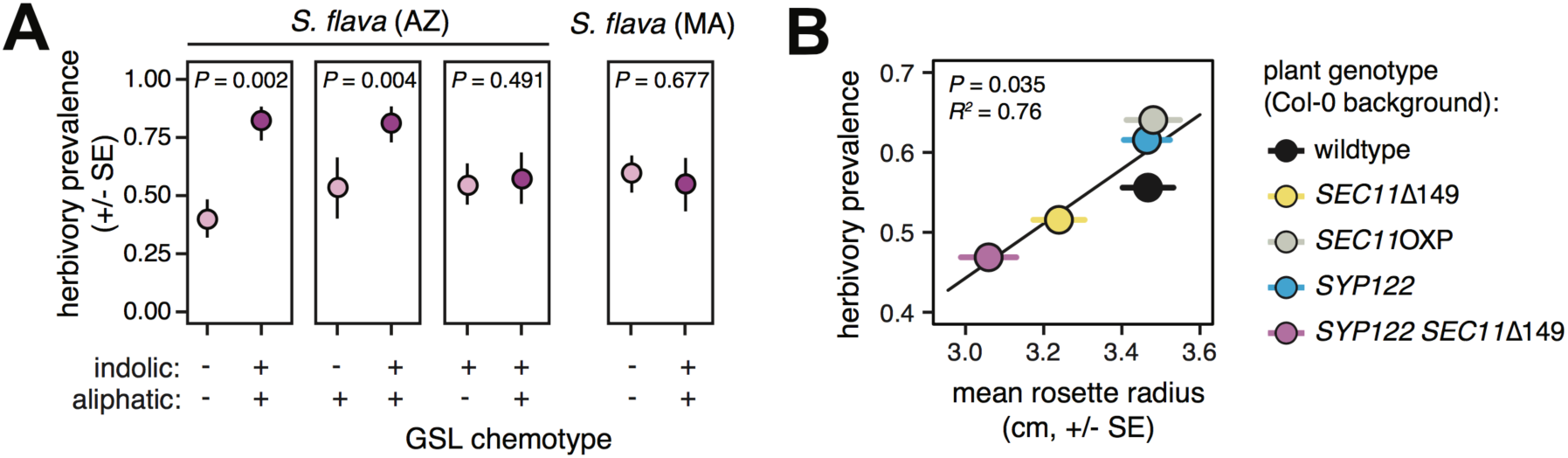
Host choice assays using *Arabidopsis* mutants verify that plant GSL profile and size mediate herbivory by *S. flava.* **(A)** Proportion of plants punctured in pairwise choice tests between *Arabidopsis* Col-0 plants with wild-type GSL profiles and mutant genotypes lacking aliphatic GSLs, indolic GSLs, or both; see methods for a description of paired genotypes in each experiment. *S. flava* AZ flies preferred plants with indolic GSLs whereas *S. flava* (MA) flies did not differentiate between plants with and without GSLs, consistent with genetic correlation analyses and GWAS. Significance and effect sizes were estimated using a binomial mixed model with replicate as a random effect and plant genotype as a fixed effect. *N* = 20 plants per replicate and 4-5 replicates per experiment. **(B)** Proportion of plants punctured in five-way choice tests, in which *S. flava* AZ flies were simultaneously presented with four *Arabidopsis* mutants varying in size due to engineered variation in expression or protein function of *SEC11* (a candidate gene at PBSL; mutants described in ^51^) or wild-type plants. Herbivory prevalence and plant size were positively correlated, showing that variation in *SEC11* expression and function can yield variation in herbivory phenotypes, consistent with GWAS results. *N* = 2 replicates of 160 plants each.

#### Effect of plant size

Before using genetic mutant plants to investigate the role of plant size on herbivory, we first conducted genetic fine-mapping with imputed *Arabidopsis* wholegenome sequences ^49^ to identify candidate genes at PBSL. Associations between PBSL genotype and plant size or herbivory phenotypes peaked in two adjacent genes: *SEC11* (AT1G12360) and *UV-B RESISTANCE2* (*UVR2*, AT1G12370) (Fig. S6). Loss of function mutations at *SEC11* and *UVR2* have been shown to reduce plant size ^50, 51^, so these genes are both strong candidates to explain phenotypic variation at PBSL (Supplementary Result 1). However, we were unable to experimentally investigate the link between *UVR2*-mediated plant size phenotypes and herbivory because reduced plant size in *UVR2* loss of function mutants is coupled with extreme necrosis ^50^. We therefore used mutant genotypes to investigate the hypothesis that *SEC11* function affects herbivory, but we acknowledge that effect of *UVR2* on herbivory also warrants investigation.

We measured susceptibility to herbivory for a suite of *SEC11* mutant genotypes ^51^ that varied in rosette size as a result of altered *SEC11* function or expression level. Herbivory prevalence was positively correlated with rosette size across this suite of mutants (Fig. 4B). Previous research using the same suite of *SEC11* mutants as our herbivory assays discovered that differences in plant size among *SEC11* mutants arise through differences in cell size ^51^. Suggestively, we found that plants in our mapping population with the derived PBSL genotype (Fig. S7), which had lower prevalence of herbivory and smaller rosettes (Table 2 & Fig. 5A), also had smaller leaf epidermal cells (Fig. 5B). We therefore propose a possible model where *SEC11-*mediated effects on cell size, through their effects on overall plant size, affect plant apparency and consequently the prevalence of herbivory damage.

**Figure 5.**
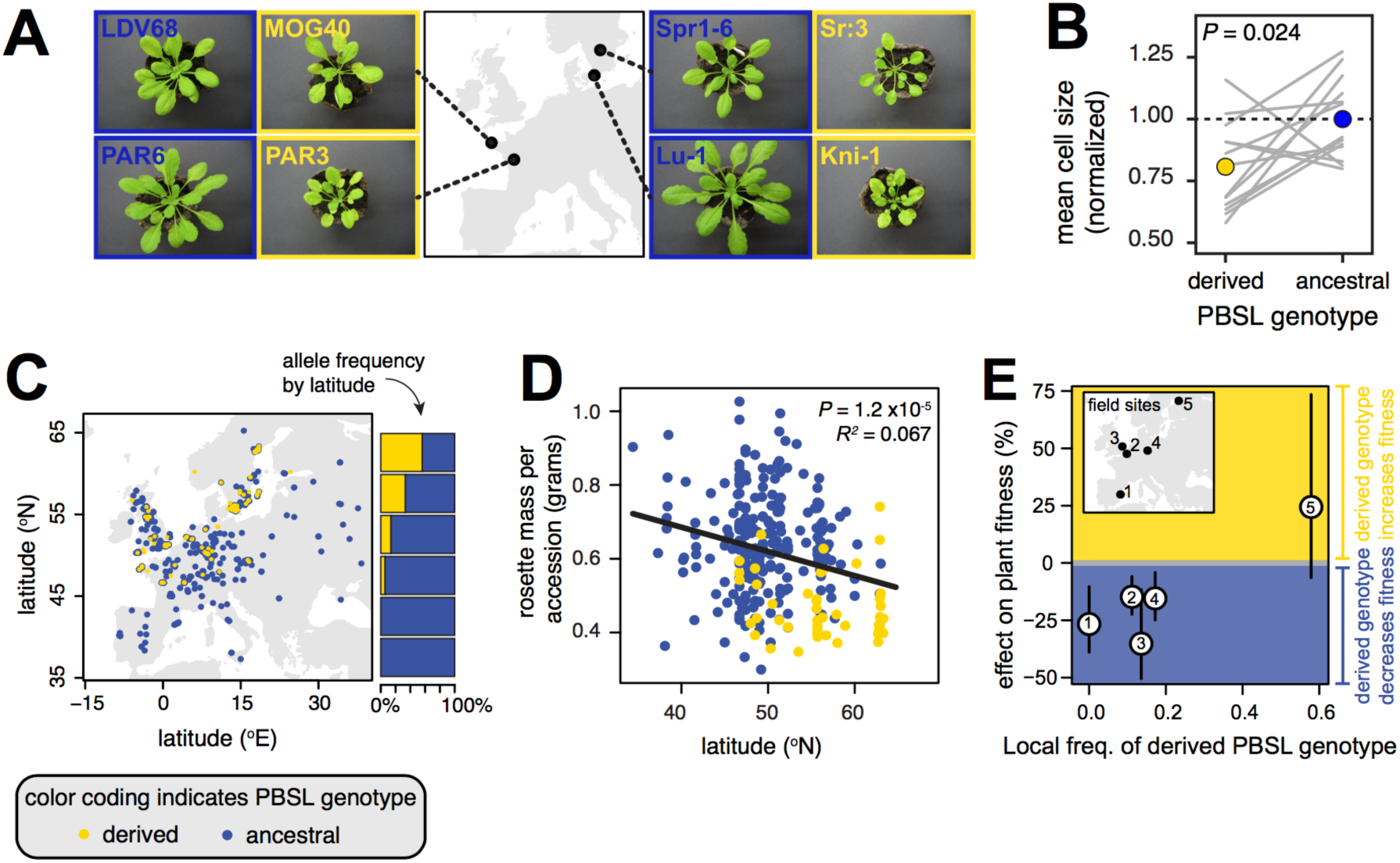
A latitudinal cline in allele frequency at PBSL can explain a cline in rosette mass among natural *Arabidopsis* accessions across Europe. **(A)** Examples of size differences among geographically paired accessions of the same age (6 weeks) differing by PBSL genotype. Determination of the derived and ancestral PBSL genotypes relied on phylogenetic analysis (see Fig. S7). **(B)** Variation in mean area of leaf upper epidermal cells among geographically paired accessions (*N* = 26 total accessions) differing by PBSL genotype. Gray lines indicate accession pairs; colored circles indicate genotype means. **(C)** PBSL genotype exhibits a strong latitudinal cline in Europe, with the globally minor allele reaching >50% frequency in northern Sweden. **(D)** Plant rosette mass is negatively correlated with latitude in *Arabidopsis* across Europe. The correlation between mass and latitude remains after accounting for population structure (*P =* 0.009), but not when both latitude and PBSL genotype are included as predictive variables in the model (*P*_latitude_ = 0.54; *P*_PBSLgenotype_= 2.9 x 10^-9^). This indicates that PBSL genotype is likely the major genetic cause of the cline in rosette mass. **(E)** The locally common PBSL genotype is associated with higher seed set (a proxy for lifetime individual fitness) in a joint analysis of five published common garden experiments ^17,56^ in the field in Europe (Stouffer’s *Z*-test, *P* = 0.03), consistent with PBSL allele frequency being driven by local adaptation. Fitness in the field was previously quantified for accessions originating throughout the global range of *Arabidopsis* (mean *N* =186 accessions per experiment), and we inferred the effect of PBSL genotype on fitness using a mixed model with an identity-by-state kinship matrix as a random effect to account for genetic background. Local genotype frequency was calculated using accessions from the RegMap panel collected within 500 km of each field site.

Overall, congruence between GWA mapping, trait-based regression analyses, and assays with mutant genotypes point to substantial effects of both plant size and GSL profile on herbivory by *S. flava.* They also suggest that a lower incidence of herbivory might arise through reduced apparency due to smaller epidermal cell size, but further experiments are needed to functionally validate the causal polymorphism and underlying gene(s) through which variation at PBSL affects plant phenotypes.

### Natural selection shapes clinal geographic variation at a locus jointly affecting plant size and herbivory

Given the pervasive effect of PBSL on plant size and susceptibility to herbivory, we hypothesized that PBSL may play a role in local adaptation in *Arabidopsis.* We first searched for population genetic patterns indicative of positive natural selection using data from the *Arabidopsis* RegMap panel ^52^. We found no evidence for near-complete global selective sweeps, partial selective sweeps, or geographic differentiation among regional populations (Table S2). However, PBSL genotype was strongly correlated with latitude of origin in European accessions (Table S3). The cline reflects an increase in the frequency of the derived PBSL genotype (Fig. S7) in accessions from northern latitudes (Fig. 5C).

Strikingly, the allelic cline at PBSL mirrored a latitudinal cline in *Arabidopsis* size that was reported previously ^53^ and recovered in our study (Fig. 5D). Specifically, latitude and plant biomass were negatively correlated (*P* =1.2 x 10^-5^, *R^2^* = 0.064). This relationship remained significant after accounting for neutral population structure by including a matrix of relatedness among accessions in a linear mixed regression model (*P* = 0.009), which suggests that the cline is unlikely to reflect random geographic sorting of genetic variation. Genomic regions underlying clinal variation in *Arabidopsis* size across Europe have not been identified, so we next tested if allelic variation at the PBSL locus contributes to this cline. When PBSL genotype was also included in the regression model, PBSL genotype was associated with plant size (*P* = 2.9x10^-9^) but latitude was not (*P* = 0.54). Although plant size ^54^, like human height ^55^, is a polygenic trait, our analyses indicate that putatively adaptive latitudinal variation in plant size in *Arabidopsis* might be caused predominately by allelic variation at PBSL.

To more directly test if the latitudinal cline at PBSL is driven by local adaptation, we quantified the effect of PBSL genotype on fitness in common garden datasets ^17,56^ from five European field sites. To account for effects of genetic background, we employed a mixed model with a matrix of relatedness among accessions as a random effect. Accessions with the locally common PBSL genotype had higher seed set (a component of lifetime fitness) than accessions with the alternate genotype (mean effect size = 23% increased seed set, 95% CI = 6-40%, meta-analysis *P* = 0.03, Fig. 5E). We cannot exclude the possibility that this pattern is driven only by a fitness cost of the derived PBSL allele in central and southern Europe because only a single garden experiment was conducted in northern Europe. Nonetheless, the presence of a strong geographic cline at PBSL, combined with a predicted fitness benefit of the locally common allele in all five field sites, suggests geographic variation at PBSL is shaped by local adaptation to latitudinally varying selective agents.

Intriguingly, the pattern of clinal variation at PBSL is discordant with latitudinal patterns in the strength of herbivory. The proportion of plant biomass consumed by herbivores is inversely correlated with latitude in a recent meta-analysis of many plant species ^57^, suggesting pressure to evade herbivory might be stronger at lower latitudes, yet the PBSL allele associated with smaller plant size and lower prevalence of herbivory in our experiments is most abundant at higher latitudes. One possible explanation for this pattern is that selective agents other than herbivory have shaped the geographic distribution of allelic variation at PBSL. Two abiotic selective agents that vary with latitude—temperature and UV radiation ^58,59^—are promising candidates that may shape the cline in PBSL allele frequency (Table S4), although interactions with herbivores should not be discounted (Supplementary Result 2). Even if clinal variation at PBSL is not driven by herbivory *per se*, it is reasonable to predict that variation at this locus might still impact the prevalence of herbivory and the fitness cost imposed by herbivores in natural populations.

### Loci mediating susceptibility to herbivory are polymorphic within and among natural populations

Our GWA mapping population was composed of *Arabidopsis* accessions originating throughout Europe, which enabled a deep breadth of species-wide genetic variation to be interrogated for loci influencing herbivory. The ecological effects of variation at these loci, however, depends in part on the extent to which they are polymorphic within stands of plants in natural populations ^15^. If a locus that mediates susceptibility to an herbivore is not polymorphic within a population, it will have no effect on which individuals in that population best evade attack by herbivores. On the other hand, if a locus is polymorphic within a population, it can impact herbivore foraging behavior, cause fitness differences among individual plants, and facilitate rapid local adaptation in response to herbivory.

We first profiled natural variation at herbivory-associated loci in 15 European *Arabidopsis* populations, each covering an area of approximately 1 km^2^ or less, using genotyped accessions from the RegMap panel ^52^ (Fig. 6A). At each locus, we focused on the top SNP that affects herbivory and GSL profile (GS-OH, AOP2/AOP3, and MAM1 loci) or that affects herbivory and plant size (PBSL). The majority of populations were polymorphic at PBSL (9 of 13 populations after excluding northern Sweden, where *Arabidopsis* exhibits strong local population structure ^60^) and at one or more GSL-associated loci (11 of 13 populations; Fig. 6B).

**Figure 6.**
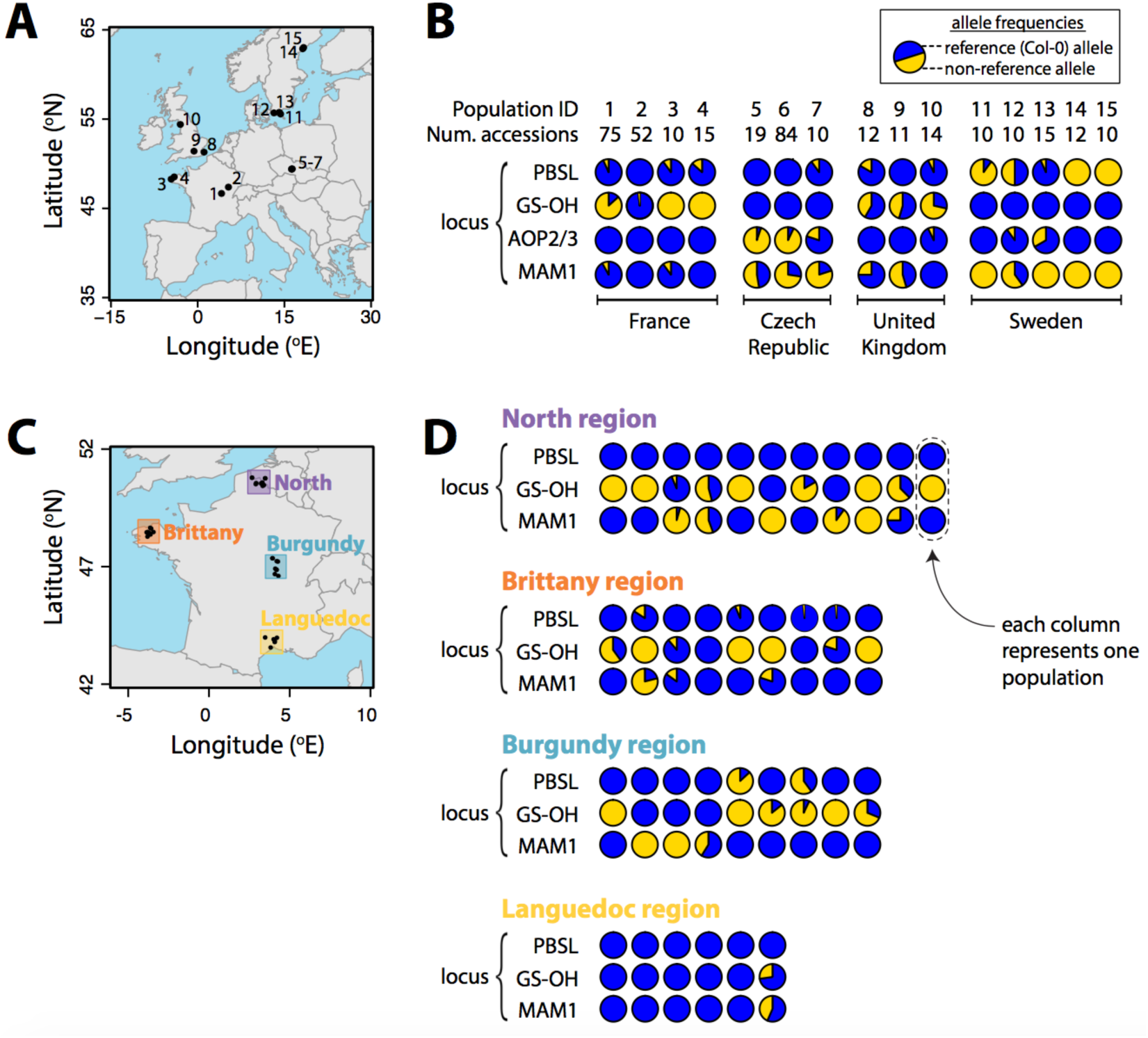
Loci mediating herbivory by *S. flava* are polymorphic within natural *Arabidopsis* populations. **(A)** Locations of 15 populations with at least 10 accessions genotyped in the RegMap panel ^52^. A population was defined as a set of accessions for which GPS collection localities differed by no more than one hundredth of a degree in latitude or longitude, corresponding to an area of approximately 1 km^2^ or less. **(B)** Allele frequencies in European populations for the top SNPs jointly associated with herbivory and either plant size (PBSL: chromosome 1, position 4202898) or GSL profile (GS-OH: chr. 2, pos. 10823986; AOP2/AOP3: chr. 4, pos. 1377499; MAM1: chr. 5, pos. 7676528) in our assays. Population ID refers to labeled locations in panel A. **(C)** Locations of populations from four regions of France that were genotyped using pooled genome resequencing. Plants within each population were collected across an area of less than 0.5 km^2^. At least 10 individual plants were collected (mean *N* =18 plants) from each population. **(D)** Allele frequencies in French populations from panel C, ordered within regions by increasing latitude, for the top SNPs associated with herbivory and either plant size (PBSL: chromosome 1, position 4202898) or GSL profile (GS-OH: chr. 2, pos. 10823986; MAM1: chr. 5, pos. 7676528) in our laboratory assays. Each locus was sequenced a mean of 17 times per population (minimum = 6x, maximum = 35x).

To interrogate variation at a finer spatial scale, we measured allele frequencies across each of 35 narrowly delimited *Arabidopsis* populations in France, each covering 0.5 km^2^ or less, using a pooled genome sequencing approach (Fig. 6C). We focused on the same SNP at each locus as described above, except the AOP2/AOP3 locus was excluded because genotypes could not be confidently inferred using our approach. Herbivory-associated SNPs again exhibited considerable variation within and among populations (Fig. 6D). The focal SNPs at GS-OH and MAM were polymorphic in 31% and 26% of sampled populations, respectively, and within all four geographic regions sampled within France. Both of these loci exhibited cases where opposite alleles predominated in populations separated by small geographic distances. The focal SNP at PBSL was polymorphic in 13% of the sampled populations and within two of four regions, even though the derived allele at PBSL was rare across the sampled French populations collectively relative to northern Europe.

In both our analyses, the number of polymorphic populations could increase with greater sequencing and sampling depth, so our results represent a minimum estimate of the extent of within-population polymorphism.

## DISCUSSION

Identifying which plant traits most strongly shape the outcome of plant-herbivore interactions remains contentious ^8,9,61^. Using GWAS to identify genetic polymorphisms affecting susceptibility to herbivory—and then determining which biochemical, physiological, morphological, and developmental traits are influenced by these same loci— offers a powerful and relatively unbiased approach to identify traits mediating plant-herbivore interactions. Experimental and population genomic studies of these polymorphisms can then shed light on how herbivory-related traits evolve in natural populations. Here, we employed GWA mapping to uncover plant traits and underlying genetic polymorphisms that shape interactions between *Scaptomyza* and *Arabidopsis*. Importantly, our experimental assays allowed herbivores to forage among a phenotypically diverse assemblage of plant genotypes, and therefore captured the importance of evasion and resistance strategies employed by plants and their effects on herbivore host choice and feeding behavior. For plants such as *Arabidopsis*, which frequently occur at densities of tens to hundreds of individuals per square meter in nature ^62-64^, traits that shape patterns of herbivore foraging among genotypes are likely to be ecologically and evolutionarily important.

Our approach to model variation in susceptibility to herbivory among plant genotypes relied on simple statistical methods to overcome two challenges. The first challenge arises because susceptibility to herbivory is a complex trait ^10^. Complex trait phenotypes—those that are jointly determined by phenotypes of a number of simpler traits—are often influenced by small effects of many genes collectively. Reducing complex traits into simpler component parts, each controlled by a smaller number of loci, can increase the power to detect genetic variants affecting those traits ^10^. We decomposed herbivory into two mechanistically distinct components, reflecting either host choice (herbivory prevalence) or feeding rate (herbivory intensity), that each had a significant and distinct genetic basis. The second challenge arises due to stochastic spatial aggregation that is typical of host-enemy interactions as enemies move across a landscape ^41^. We employed a spatially explicit statistical model to minimize error introduced into our phenotypic measurements due to stochastic spatial aggregation. These approaches, which could be applied to studies of other host-enemy systems, enabled the detection of known and novel loci influencing susceptibility to herbivory in *Arabidopsis.*

Allowing herbivores to move freely among plants in large enclosures during feeding assays revealed that plant size had a substantial effect on the probability that a plant would be attacked. Strikingly, this effect was due in part to a single locus with a large effect on the probability of attack, but no effect on feeding rate on attacked plants. This pattern is consistent with the core premise of plant apparency theory, which predicts that larger plants will be attacked more often by herbivores because they are encountered more frequently ^20,61^. The effect of plant apparency on herbivory is well supported in trait-based ecological studies (e.g., ^35-37^). However, the discovery of loci jointly affecting plant size and the probability of attack by herbivores has remained elusive in genetic mapping studies, likely because these studies typically place herbivores directly onto plants rather than allowing them to forage. Consequently, results of genetic mapping studies have traditionally emphasized the importance of chemical defenses for resistance to herbivory. Our results show that allowing herbivore foraging among plant genotypes during genetic mapping studies points to both plant size and chemical defenses as major determinants of herbivory, at least for *Arabidopsis* and *Scaptomyza*, and thus reduces discrepancies between trait-based ecological studies and genetic mapping studies. Extending this approach to more natural field habitats, however, will be key to further reducing differences between genetic mapping studies and trait-based ecological studies. For example, the effect of plant apparency on variation in herbivory among *Arabidopsis* genotypes might increase as herbivore foraging extends over larger spatial scales, or decrease as mustard-feeding herbivores encounter a range of interspecific variation in size in the Brassicaceae that is far greater than the range of variation within *Arabidopsis*.

Although our genetic mapping approach found that both plant size and chemical defenses mediate herbivory by *S. flava*, our results also suggest that the effects of plant size and chemical defenses may differ in their specificity. In the case of GSL-mediated effects, the outcome of attack by *S. flava* depended simultaneously on both the host plant’s chemotype (and genotype at the underlying GSL-associated loci) and the herbivorous insect’s geographic provenance (AZ vs. MA). In other words, genotype_plant_ × genotype_herbivore_ interactions drove chemical-mediated effects on host choice and feeding rate. Such G × G interactions frequently influence the outcome of host-parasite interactions ^65^, but are not well understood in the context of herbivory. Conversely, we did not find evidence for G_plant_ × G_herbivore_ interactions in the context of plant size. Instead, we found a strong G_plant_ effect on the probability that a plant would be attacked: larger plants were more likely to be attacked regardless of the herbivorous insect’s geographic provenance. An intriguing general hypothesis arising from these results—which will be testable as GWA mapping approaches are employed for other plant and herbivore species—is that effects of chemical defenses might frequently be driven by G_plant_ × G_herbivore_ interactions, while effects of plant size may be less dependent on herbivore genotype.

Our finding that plant size- and GSL-associated loci mediating interactions with herbivores are polymorphic within and among natural populations, at a finer spatial scale (less than 1 km^2^) than previously examined in depth in *A. thaliana*, suggests these loci could shape the feeding behavior of herbivores foraging within stands of *Arabidopsis* in the field. Previous studies have documented that GSL profiles and GSL-modifying QTL vary within populations of *A. lyrata*, an outcrossing relative of *A. thaliana*, and that this variation affects plant fitness and resistance to herbivory ^66, 67^. Our analyses suggest that genes determining accumulation of different GSL molecule structures are polymorphic within many populations of *A. thaliana* as well, despite the reduction in genetic diversity that results from a high rate of selfing in this species ^68^. Further genomic analyses are needed to determine if loci mediating GSL phenotypes harbor more genetic variation than loci across the genome that do not affect plant phenotypes or fitness, as predicted if spatial and temporal fluctuation in natural selection maintains adaptive genetic variation at loci shaping GSL profiles ^16^.

At a broader spatial scale, population genetic analyses revealed strong geographic clines in Europe for alleles at GSL-associated loci (e.g., ^6,17^) and plant size-associated loci (this study) that mediate herbivory by *S. flava.* In the case of GSL-associated SNPs, this pattern has been shown to have arisen, at least partially, in response to selective pressures exerted by local herbivore communities ^6,17^. Intriguingly, however, clinal variation at the plant size-associated locus discovered in the current study was discordant with the general latitudinal trend in the intensity of herbivory. Specifically, the allele associated with evasion of herbivory was more frequent in higher latitudes, whereas the strength of herbivory is generally greater in lower latitude regions for plants in temperate regions ^57^. This discordance might arise if other agents known to exert selection on plant size and growth, such as climate or biotic competition, exert stronger pressure than herbivores on loci affecting plant size. Such a scenario is consistent with the hypothesis that secondary chemicals have evolved major roles in defense against herbivores partly because their evolution is less constrained than other traits, such as plant growth and morphology ^9^. In the future, a combination of field experiments and population genetic studies could reveal whether there is commonly a geographic discordance between the strength of herbivory and the distribution of alleles involved in evading herbivory through reduced size. However, even for our study, evidence for such discordance remains tentative without quantifying the strength and fitness effects of herbivory on plants with alternate PBSL genotypes in field sites across Europe.

Our study focused on polymorphisms within a single species, but the same loci may have important effects across other taxa as well. A major question following from the present study is whether natural or engineered variation at PBSL affects biomass in other species in the genus *Arabidopsis*, the family Brassicaceae, and beyond. If so, genetic manipulation of this locus through transgenic or conventional methods may lead to the ability to precisely alter plant biomass, and, possibly, affect rates of herbivory. Genes mediating interspecific differences in plant size likely explain some of the variation in herbivory across species in the family Brassicaceae, given that larger species in this family incur more herbivory damage than small species ^69^, but individual genes underlying this pattern have not yet been identified.

## Conclusion

We hope our conceptual and statistical approach will motivate future attempts to dissect genetic variation mediating host-enemy interactions that emphasize the importance of enemy behavior in heterogeneous environments. In our study, such an approach highlighted how canonical defense traits (specifically, secondary chemicals) but also traits other than canonical defenses (such as plant size and perhaps phenology) influence susceptibility to herbivory, consistent with ecological theory and observations ^9^. If our results are typical of plant-herbivore interactions, extending similar experimental designs to other plant and herbivore species and to more natural habitats in the field is likely to broaden the catalog of polymorphisms known to mediate plant-herbivore interactions. Given that polymorphisms affecting plant growth and physiology typically exhibit less spatiotemporal variation in allele frequency than those affecting canonical defense traits ^52^, an increased emphasis on traits other than canonical defenses could shed new light on the processes that constrain the evolution of plant defense against herbivory.

## ACKNOWLEDGMENTS

We thank Timothy Morton, Timothy O’Connor, Ingrid Peterson, and Amelia White for assistance with experimental setup or data collection and processing. We also thank Mark Beilstein, Katrina Dlugosch, and Michael Nachman for feedback on drafts of the manuscript. Research reported in this publication was supported by grants from the National Science Foundation (DEB-1256758 to NKW and DEB-1405966 to ADG and NKW), the John Templeton Foundation (award 41855 to NKW), the National Institute of General Medical Sciences of the National Institutes of Health (R35GM119816 to NKW and GM083068 to JB) the University of California, Berkeley (laboratory setup funds to NKW), the Région Midi-Pyrénées (CLIMARES project to FR), the LABEX TULIP (ANR-10-LABX-41 and ANR-11-IDEX-0002-02 to FR), and the Commission Franco-Américaine Fulbright (fellowship for research scholars to FR). The content is solely the responsibility of the authors and does not necessarily represent the official views of the National Science Foundation or the National Institutes of Health.

## AUTHOR CONTRIBUTIONS

ADG, BB, JB and NKW conceived and designed the project. ADG, MJF, SCG, CB, JG, ERL, CGM, HSP, SCR, and FR performed the experiments and collected or processed data. ADG carried out the analyses and wrote the paper with feedback from BB, MJF, SCG, JB and NKW. All authors contributed to revisions of the paper.

## METHODS

### Plant and Fly Populations

The *Arabidopsis thaliana* (*Arabidopsis* hereafter) mapping population for GWAS consisted of 288 natural accessions chosen to encompass phenotypic and genetic diversity across the native range within Europe. The population included the core set of accessions from Atwell et al. ^42^, but we excluded most accessions of non-European origin. Additional accessions from the RegMap panel ^52^ were randomly added to include an even sampling of accessions from Eastern and Western European populations.

Laboratory *S. flava* populations were founded from wild-collected larvae from two sites in North America: Belmont, MA, USA (see ^38, 40, 70^ for details on the natural history of this population in relation to *Arabidopsis*) and Flagstaff, AZ, USA. Both populations were founded within one year, or approximately 15 generations, prior to the initiation of experiments and subsequently maintained on wild-type *Arabidopsis* Col-0 plants. Flies were not specifically inbred because this species tolerates inbreeding poorly.

Interactions between *S. flava* and *Arabidopsis* populations in this study have not been shaped by a history of tight, pairwise co-evolution because *S. flava* feeds on a variety of species in the Brassicales ^38^, *Arabidopsis* is attacked by many herbivore species ^71^, and we exposed North American populations of *S. flava* to *Arabidopsis* accessions primarily from Europe. Such a scenario, where evolution of both the plant and herbivore has been shaped by interactions with many different species, is typical in temperate regions ^72^. Accordingly, the patterns we observed for *S. flava* and *Arabidopsis* may be broadly relevant.

### Plant and Fly Rearing

Prior to planting, seeds were sterilized in 10% aqueous bleach and cold-stratified in sterile water for 5 days at 4°C without light. Individual plants were grown in Jiffy 7 peat pellets in 6 cm wide divided cells under a 16/8 hr, 21/15° C light/dark cycle with approximately 50% humidity in climate controlled rooms at the University of Arizona. The location of each plant was randomly assigned. To minimize positional effects, cells were rotated within trays, and trays were rotated across shelves, 2-3 times per week. Light was maintained at 4000-5000 lux, near the optimal intensity for *Arabidopsis* growth ^73^, using F40T12 Cool White Supreme fluorescent tube lights (Philips Lighting, Somerset, NJ). Beginning 2 weeks post-germination, plants were bottom-watered with the fertilizer mixture described in Brachi et al. ^17^. For the experiments using *SEC11* mutants, all plants were misted until dripping with 30 uM Dex in distilled water once per 48 hrs to induce expression of *SEC11* constructs. All experiments were conducted 42-46 days after planting, using identically aged plants within each experimental replicate. Individual plants that had begun flowering were included in the experiment but excluded from further analysis.

Flies used in experiments were raised to adulthood and mated on wild-type *Arabidopsis* Col-0 plants. *S. flava* has not been successfully reared on an artificial diet, and the *S. flava* AZ population had very low oviposition and feeding rates on *myb28 myb29 cyp79b2 cyp79b3* plants ^48^ deficient in glucosinolates, so we could not obtain naïve flies that had not fed on glucosinolates prior to the experiments. Individuals used in experiments were briefly (~3-5 days) kept in vented 50 mL plastic centrifuge tubes following adult eclosion, where they were provisioned with cotton wool soaked in 10% honey solution in water. Flies were subsequently released into 33 x 33 x 33 cm white mesh cages with clear plastic tops (Live Monarch Foundation, Boca Raton, FL), where they were provisioned with Col-0 wild-type plants and cotton wool soaked in 10% honey solution, and were collected for use in experiments 1-2 weeks after release into cages. Flies were only anesthetized with CO_2_ once, during the transfer to plastic tubes following adult eclosion.

### Herbivory Assays

For all assays, fly feeding punctures per plant were counted by eye using a Zeiss Stemi 2000C microscope (Zeiss Microscopy, Jena, Germany) under 6.5x magnification. Assays were conducted at 21°C in a climate controlled room at the University of Arizona.

#### Feeding preference assays (choice tests) with natural accessions

For each replicate, 288 plants (one of each *Arabidopsis* accession in the mapping population) were randomly arrayed at 5.5 cm intervals along a grid in a 183 x 61 x 61 cm cage with white mesh sides and a clear plastic top (Live Monarch Foundation, Boca Raton, FL). Fifty gravid female flies and 50 male flies were released into the center of the cage and allowed to feed for 24 hours on a 16/8 hr light/dark cycle. The cage was rotated 180° once after 8 hours of daylight to reduce positional effects. The experiment was replicated four times with *S. flava* AZ flies and two times with *S. flava* MA flies. Individual flies were not reused across experiments.

#### Feeding preference assays (no choice tests) with natural accessions

For a subset of the full mapping population (N = 102 accessions), plants were individually placed in a Magenta tissue culture box (7.6 x 7.6 x 10.2 cm) with white cloth mesh secured on the top. A single, randomly-assigned, gravid female fly was released into each cage and removed after 70 minutes. The experiment was conducted with *S. flava* AZ flies only and replicated twice per accession across two batches. Each fly was re-used 3-4 times within a batch, and separate flies were used for each batch to minimize effects of past experience. Non-damaged plants (< 5% of trials) were re-assayed with new flies. In all cases, plants were damaged in the repeated trials, suggesting initial measurements of 0 punctures were due to inactive flies or genetic variation among flies, so these initial measurements were discarded and re-measurements were retained.

#### Feeding preference assays (choice tests) with mutant genotypes

Assays using mutant plants varying in size or GSL profile were conducted to complement analyses using natural variation. A suite of mutants varying in expression or function of *SEC11* were used to test if *SEC11*-mediated size variation affects prevalence of feeding punctures from *S. flava* AZ flies. Five genotypes in the Col-0 genetic background, previously described in Karnik et al. ^51^, were used: wildtype, *SEC11*Δ149 (a *SEC11* reduction-of-function mutant with reduced rosette size), *SEC11* OXP (an overexpression mutant with increased rosette size), *SYP122 SEC11*Δ149 (exhibits stronger dwarfism than the *SEC11*Δ149 mutant due to interaction with *SYP122*), and *SYP122* (no visible change in size relative to wild-type). Three independently transformed lines were used for mutants containing the *SEC11*Δ149 or *SEC11* OXP constructs. The assay was conducted as described in “Feeding preference assays (choice tests) with natural accessions*”* with the following exceptions: 155 plants were used per replicate (N = 31 plants per genotype), 27 female and 27 male flies were used to maintain the same plant:fly ratio, and two replicates were conducted.

To test if total GSL affect puncture prevalence from *S. flava* MA and *S. flava* AZ herbivory, pairwise choice tests were conducted with wild-type Col-0 plants and mutants deficient in all GSL (*myb28 myb29 cyp79b2 cyp79b3*, ^48^; hereafter GKO for Glucosinolate Knock Out). The assay was conducted as described in “Feeding preference assays (choice tests) with natural accessions*”* with the following exceptions: 8 gravid female flies were placed in a 31 x 31 x 31 cm white mesh cage with 10 wild-type plants and 10 GKO plants for 8 hours, and tests were replicated four times for each fly population. Additional choice tests were performed with the *S. flava* AZ population to test if indolic and/or aliphatic GSL affect feeding preference. In the first test, flies were released in cages with mutants deficient in indolic GKO (*cyp79b2 cyp79b3*, ^47^; hereafter iGKO for indolic Glucosinolate Knock Out) and *pad3-1* control plants (N = 5 replicates). *pad3-1* mutants were used rather than wildtype as controls because they produce GSL but lack camalexins, a defensive compound also absent in GKO and iGKO mutants ^74,75^. In the second test, flies were released in cages with mutants deficient in aliphatic GKO (*myb28 myb29*, ^46^; hereafter aGKO for aliphatic Glucosinolate Knock Out) and wildtype control plants (N = 4 replicates). iGKO mutants exhibit reduced indole acetic acid levels and growth defects consistent with partial auxin deficiency at high temperatures ^47^, but we did not observe substantial size differences among wild-type plants and the *pad3-1*, GKO, iGKO, and aGKO mutants in the growing environment maintained during this study.

### Rosette Size and Leaf Cell Size Assays

Three plants per each accession were grown in randomly assigned positions as described in “Plant Rearing”. Plants were harvested between 43 and 45 days after planting (N = 270 plants per day) by clipping at the lower base of the rosette, and stalks were removed from any flowering plants. Rosettes were immediately photographed and weighed on a CPA225D microbalance (Sartorius, Gottingen, Germany). Following drying at 60°C for 72 hrs, plants were weighed to determine dry mass. Rosette diameter was quantified in imageJ ^76^.

Leaf upper epidermal cell size was measured for a subset of the mapping population consisting of 15 pairs of accessions differing in genotype for the SNP position 4202898 of chromosome 1 (e.g., each of the two PBSL alleles). To construct the reduced mapping population, 15 accessions with the minor allele were randomly chosen, and each accession was paired with the geographically closest accession with the major allele. For each plant (N = 2-3 plants per accession), leaf 4 was harvested 42 days after planting and the epidermal surface was painted with clear nail polish. After air drying, the mold was removed by pressing the leaf against clear tape and peeling away leaf material with a forceps, and the mold was secured onto a microscope slide by pressing down the surrounding tape. Cell molds were photographed using an EOS T3i camera (Canon Inc., Tokyo, Japan) on a Leitz Laborlux D microscope (Leica Microsystems, Wetzlar, Germany). Area was calculated by a single observer for two randomly selected cells per leaf photograph, excluding small stomata-associated cells, through manual tracing using ImageJ ^76^.

### Glucosinolate Assays

Abundance of 22 aliphatic GSL compounds was previously quantified in 3 week-old rosettes of 595 accessions from the RegMap panel and reported by Brachi et al. ^17^. New measurements, generated from the same plant tissue and HPLC runs as those in Brachi et al. ^17^, were included in this study for two indolic glucosinolates: indol-3-yl-methyl (I3M; *m/z* = 446.2, retention time = 153 s) and methoxy-indol-3-yl-methyl (MI3M; *m/z* = 477.06, retention time = 260 s). Two liquid chromatography-MS/MS runs with low indolic GSL detection were discarded from further analyses of indolic or aliphatic GSLs profiles, leading to slight reduction in replicate number for some accessions compared to Brachi et al. ^17^.

### Estimation of Genetic Effects and Heritability

We tested whether plant accession ID (i.e., plant genotype) had a significant effect on five trait classes, including two classes of herbivory traits: (1) herbivory prevalence, a binary trait indicating whether plants were damaged or undamaged, and (2) herbivory intensity, the puncture number for damaged plants. Phenotypes were modeled using GLMMs implemented in R (packages: *glmmADMB & lme4* ^77^) with accession ID as a random effect, and the statistical significance of accession ID was determined using a likelihood ratio test. Random and fixed effects, error distribution, transformations, and the unit scale of the conditional modes or BLUPs are as follows:

| Trait | Model effects<br>(R = random;<br>F = fixed) | Error<br>distribution | Pre-modeling<br>transformation | Scale of conditional<br>modes or BLUPs |
| --- | --- | --- | --- | --- |
| herbivory prevalence | R: ID, replicate;<br>F: positional effect | binomial | none | log(odds ratio) |
| herbivory intensity | R: ID, replicate,<br>F: positional effect | neg. binomial | none | log(puncture count) |
| rosette mass | R: ID, replicate,<br>growth shelf<br>position, plant age | normal | log(x+1) | log(mass + 1) |
| rosette diameter* | R: ID, replicate,<br>growth shelf<br>position, plant age | normal | $\pi(\frac{1}{2}x)^2$ | circular area |
| GSL profile | R: ID, replicate | normal | log(x+1) | log(ion count/mg + 1) |

\* Rosette diameter per plant was transformed to circular area for modeling remove a right-tailed skew in the distribution of residuals

We used two methods to account for aggregation of herbivory damage (i.e., many host individuals are lightly attacked and few host individuals are heavily attacked), a typical pattern in host-parasite interactions ^41^, in GLMMs. To account for individual-level aggregation, we modeled herbivory intensity per plant with a negative binomial error distribution. To account for aggregation within regions of experimental cages, the mean number of feeding punctures on immediately adjacent plants (i.e., in a 3 x 3 plant grid centered on the focal plant) was included as a fixed effect with seven bins containing roughly equal numbers of focal plants: < 10,10-20, 20-30, 30-40,40-55, 55-75, and > 75 punctures. Bins were used to ensure our results were not heavily biased by a non-linear relationship between herbivory damage on the focal plant and neighboring plants. Plants at cage edges had fewer neighbors but were still included. Individual plants were excluded if all adjacent plants were undamaged; this resulted in removal of ~1% of plants from the dataset.

Broad-sense heritability was estimated from fitted GLMMs as V_G_ / (V_G_ + V_E_), with V_G_ estimated as the variance for the random effect of accession ID and V_E_ estimated as the residual variance of each model.

### GWA Mapping

GWA mapping was conducted using the Accelerated Mixed Model (AMM, ^78^), a modification of EMMAX ^44^ where exact *P*-values are estimated for the 200 SNPs showing the strongest associations. Accession phenotypes estimated as conditional modes from GLMMs were used for puncture prevalence, puncture number, rosette mass, and rosette circular area. Because an overrepresentation of accessions with values of 0 for some GSL compounds could bias GLMM estimates of conditional modes, median log-transformed ion counts per mg of lyophilized plant tissue were used for each of 22 aliphatic and two indolic GSL compounds. Accession genotypes were obtained from the 250k SNP array from the RegMap panel ^52^. An identity-by-state kinship matrix, calculated using all SNPs in the analysis, was included in all analyses as a random effect to account for genetic relatedness among accessions. Pseudo-heritability, the proportion of phenotypic variance among accessions explained by pairwise relatedness, was estimated as described by Kang et al. ^44^. All GWA mapping and pseudo-heritability analyses were implemented using PyGWAS (https://github.com/timeu/PyGWAS).

To test for a shared genetic basis among heritable variation in susceptibility to herbivory and either plant size or GSL variation, GWAS for traits within each of three broad plant phenotypes—rosette size (wet mass, dry mass, and circular area), aliphatic GSL profile (22 total compounds), and indolic GSL profile (I3M- and MI3M-GSL)—were combined by taking the top *P-*value among all individual GWAS at each SNP. SNPs with a minor allele frequency (MAF) less than 10% in the mapping population were excluded from further analysis for all herbivory phenotypes; for plant size and GSL phenotypes, which generally exhibited higher heritability and were measured across larger mapping populations, SNPs with MAF < 5% were excluded. The enrichment of shared SNPs in the upper tails of the GWAS *P-*value distribution for each herbivory trait and either rosette size, aliphatic GSL profile, or indolic GSL profile was quantified using the equation 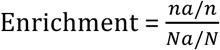, where *n* is the number SNPs in the tail of the herbivory GWAS *P-*value distribution, na is the number of those SNPs that are also in the tail of the GWAS *P-*value distribution for the plant size or GSL trait being compared, *N* is the total number of SNPs included in the GWAS, and *Na* is the number of those SNPs in the tail of the GWAS *P-*value distribution for the plant size or GSL trait. Statistical significance of enrichments was calculated by comparing the observed enrichment ratio to the distribution of enrichment ratios from 1000 genomic permutations. In each permutation, the positions of all SNPs in the herbivory trait GWAS were rotated by a random interval, preserving the order of SNPs and their associated *P-*values.

### Genomic Fine-Mapping

To localize the peak association with plant size and herbivory traits at PBSL, imputed genotypes from whole-genome resequencing were obtained from the *Arabidopsis* 1001 genomes project ^49^, and GWA mapping for SNPs in the PBSL region was implemented using an Accelerated Mixed Model as described above. We also binned accessions by their genotype at the top-scoring SNP at PBSL and employed nested GWA mapping ^79^, but this approach revealed no new associations (results not reported). Functional annotations were obtained using the GWA Portal SNP Viewer tool (https://gwas.gmi.oeaw.ac.at/).

*Arabidopsis* accessions from Sweden, which have been robustly genotyped for structural variants in addition to SNPs ^60^, were used to interrogate patterns of LD at PBSL. Only directly sequenced (i.e., non-imputed) genotypes were used. LD calculations and heatmap preparation used the *genetics* ^80^ and *LDheatmap* ^81^ R packages.

To determine whether the PBSL haplotype associated with reduced plant size is ancestral or derived in *Arabidopsis*, a maximum likelihood phylogeny of the *SEC11* and *UVR2* coding sequences was constructed using RAxML HPC-Hybrid-SSE3 v7.2.8 ^82^ under the GTR+G substitution model with *Arabidopsis lyrata* as an outgroup, and midpoint-rooted intraspecific phylogeny was constructed from *SEC11* genic sequences using neighbor-joining with default settings in MEGA 6.06 ^83^. Both phylogenies used sequences from the *Arabidopsis* 1001 genomes project ^49^ that were aligned using the MAFFT v7 webserver (http://mafft.cbrc.jp/alignment/server/).

### SNP Effect Estimation

To estimate the size of the effect of PBSL genotype (chromosome 1, position 4202898) on plant size and herbivory, phenotypes were modeled using the function *lmekin* from the *coxme* package in R ^84^. The model was Y_i_ ~ SNP_i_ + k_ij_ + ε_i_, where SNP_i_ is the genotype of each accession, Y_i_ is the phenotype for each accession, k_ij_ is the random effect of the identity-by-state kinship matrix of relatedness among accessions, and ε_i_ is error unaccounted for by other model terms. The proportion of phenotypic variance explained by the SNP in this model was extracted from PyGWAS ^78^.

The same approach, using a mixed model with an identity-by-state kinship matrix as a random effect to control for genetic background, was used to estimate the effect size of PBSL genotype on plant fitness in the field. Seed set measurements (silique number or cumulative silique length) were obtained from five published common garden field experiments with genotyped accessions ^17,56^ and square root-transformed to ensure normality of residuals in the fitted models. Local genotype frequencies were calculated using all plants from the RegMap panel ^52^ originally collected within 500 km of each field site, the minimum distance yielding at least 20 local accessions per site. The probability that the locally-common PBSL genotype increased fitness across all five experiments was estimated by combining *P-*values across experiments using Stouffer’s *Z* method ^85^ for meta-analysis. We repeated the analysis using geographic distance between the origin of each accession and the field sites, rather than a kinship matrix, to control for genetic background. The significance of the effect of PBSL genotype on fitness did not differ between the two approaches, so only results from the first approach were reported.

### Phenotypic correlations

Correlations among herbivory, mass, and GSL phenotypes for each accession were modeled using LMMs implemented using *lmekin*. Models followed the equation Herbivory_i_ ~ Mass_i_ + GSL.PC1_i_+ GSL.PC2_i_+ I3M_i_ + MI3M_i_ + k_ij_ + ε_i_, where Herbivoryi is the herbivory phenotype for each accession, Massi is wet mass, GSL.PC1_i_ and GSL.PC2_i_ are the first two principal component axes encompassing variation among six major aliphatic GSL molecules (2P, 2H3B, 3B, 3HP, 3MSP, 4MSB), I3M_i_ and MI3M_i_ are the log-transformed abundances of the two indolic GSL molecules in our study, and k_ij_ is the identity-by-state kinship matrix of genetic relatedness among accessions. Phenotypes were modeled as fixed effects, and the kinship matrix was included as a random effect. Use of dry mass or diameter instead of wet mass as a proxy for plant size yielded similar results. We included principal components capturing variation in aliphatic GSL profiles, rather than the abundance of individual aliphatic GSL compounds, because single genes in the aliphatic GSL biosynthesis pathway typically have correlated effects on multiple GSL molecules ^17,86^. The kinship term was included to minimize spurious correlations that could otherwise arise among uncorrelated traits as a result of similar geographic distributions of the loci affecting each trait. The statistical significance of the correlation between each herbivory and size or GSL phenotype was assessed using Wald’s *Z* tests. Our “full models” retained all terms that were significant at a 5% false discovery rate (accounting for five terms per model and three phenotypes modeled). The proportion of variance explained by each significant predictor variable was estimated as the reduction in variance explained (*R^2^*) of the full model when that variable was dropped from the model.

Correlations among herbivory and eight individual GSL molecule abundances for each accession were modeled using LMMs implemented as above, following the equation Herbivory_i_ ~ Mass_i_ + GSL. Mol_i_+ k_ij_ + ε_i_, where GSL.Mol_i_ is the log-transformed median abundance per accession of a given GSL molecule.

### Geographic Variation in Plant Size

Variation in mass was modeled using LMMs implemented using *lmekin* following the equation Mass_i_ ~ Lat_i_ + k_ij_ + ε_i_, where Lat_i_ is the latitude of origin for each accession as a fixed effect. The kinship component was included to test if the observed cline exceeds neutral expectations due to population structure. In a subsequent model, genotype at PBSL (chromosome 1, position 4202898) was also included as a fixed effect to test if latitudinal variation in size was independent of, or potentially the result of, the latitudinal cline at PBSL. The significance of fixed effects was assessed using Wald’s *Z* tests.

### Estimates of Allele Frequencies within French Populations

*Arabidopsis* specimens were collected previously from a total of 35 populations in four regions of France (i.e. Burgundy, Brittany, Languedoc and North) ^62^. Plants were collected across a maximum area 500 m^2^ per site. DNA was extracted individually from 10-20 plants per population, using 3-week old tissue grown from seed, with the DNeasy Plant Mini Kit (Qiagen, Germantown, MD). DNA was quantified with a PicoGreen dsDNA assay kit using a 7900 qPCR machine (Applied Biosystems, Waltham, MA). An equimolar pooling of separately extracted DNA was performed for each population, resulting in a final mean amount of DNA of 5μg. DNA genome sequencing was performed at the Argonne National Laboratory on an Illumina HiSeq2000 using a paired-end read length of 2x100 pb with Illumina TruSeq DNAseq library preparation. Reads from each of the populations were mapped onto the TAIR10 Col-0 *A. thaliana* reference genome using glint (1.0.rc8; Faraut & Courcelle, http://lipm-bioinfo.toulouse.inra.fr/download/glint/) with the following parameters: a maximum of 5 mismatches on at least 80 nucleotides, keeping alignments with the best score (*glint mappe --no-lc-filtering --best-score --mmis 5 --lmin 80 --step 2*). Prior to SNP calling, we determined the mean sequencing coverage across the nuclear genome for each population (mean = 49.7x, median = 45.9x, min = 25.3x, max =102x). This information was then used to adjust SNP calling parameters. SNP calling across the genome was performed for each accession with SAMtools mpileup (v0.01019) ^87^ and VarScan (v2.3) ^88^. To ensure that SNPs included in this study were unlikely to be copy number variants or mapping artifacts, we verified for each SNP that mean coverage depth was less than 60x.

### Population Genetics

The strength of the latitudinal cline at each SNP in *Arabidopsis* was quantified using logistic regression (latitude ~ SNP genotype) in 1078 European accessions. Additional population genetic statistics (Table S2) were retrieved from Horton et al. ^52^. Partial correlation coefficients between SNP genotype and environmental variables were obtained from Hancock et al. ^89^. To test for exceptional patterns at PBSL, the genome-wide rank of a peak plant size- and herbivory-associated SNP at PBSL (chromosome 1, position 4202898) was compared to the genome-wide distribution of each population genetic statistic for SNPs with MAF > 0.1.

### Data availability

All data and scripts used for data analysis are available from the authors upon request. Deposition in a public database is in progress; when complete, this manuscript will be updated to include instructions for accessing these data. Illumina sequences generated for the 35 populations were deposited in the SRA database under accession SRP097331.

